# CoralBleachRisk-Global projections of coral bleaching risk in the 21^st^ century

**DOI:** 10.1101/2024.04.16.589829

**Authors:** Camille Mellin, Stuart C Brown, Scott F Heron, Damien A Fordham

## Abstract

Timing, duration, and severity of marine heatwaves are changing rapidly in response to anthropogenic climate change, thereby increasing the frequency of coral bleaching events. Mass coral bleaching events occur because of cumulative heat stress, which is commonly quantified through Degree Heating Weeks (DHW). Here we introduce *CoralBleachRisk*, a daily-resolution global dataset that characterises sea surface temperatures, heat stress anomalies, and the timing, duration, and magnitude of severe coral bleaching conditions from the recent past (1985) to the future (2100) under three contrasting Shared Socioeconomic Pathways. Our projections are downscaled to a 0.5° resolution (~50km), bias-corrected and validated using remotely sensed data of sea surface temperatures and a global dataset of historical coral bleaching events. An accompanying online software tool allows non-specialist users to access aggregated metrics of coral bleaching risk and generate time series projections of coral vulnerability for Earth’s coral reefs. More broadly, our dataset enables regional to global comparisons of future trends in severe coral bleaching risk and the identification of potential climate refugia for corals.

## Background & Summary

Among all marine ecosystems, coral reefs and the ecosystem services they deliver to human populations are most at risk of human-driven climate impacts^1^. This is because reef-building corals live close to their upper thermal tolerance thresholds^2^, making them particularly susceptible to mean annual anthropogenic warming of Earth’s oceans^3^. However, an even more pervasive human-driven climate threat to coral reefs is the increasing frequency and severity of acute marine heatwaves, a modern phenomenon causing recurrent mass coral bleaching events at regional scales (>1000km)^4^ and, in many cases, coral mortality. Globally, the annual risk of coral bleaching increased from 18% to 31% in the three decades to 2016, along with a 4.6-fold reduction in the return time of severe bleaching events^4^. In some severe marine heatwave events, corals have died directly from rapid-onset extreme heat^5^.

Coral bleaching − paling of the coral host − occurs when symbiotic microalgae (zooxanthellae, i.e., *Symbiodinium* spp.) leave the coral tissue after prolonged heat stress, resulting in coral mortality if cumulative heat stress is prolonged^6,7^. Risk of coral bleaching and subsequent mortality is commonly predicted using a cumulative metric of heat stress called degree heating weeks (DHW) — the sum of all positive anomalies above a maximum monthly mean (MMM) sea surface temperature (SST) threshold over a 12-week rolling window^8,9^. While there is a strong correlation between maximum annual DHW and the probability of severe coral bleaching (and also mortality), the strength of the relationship can vary spatially and temporally^10^, owing to environmental filtering resulting from past severe bleaching events: what has been termed ‘ecological memory’^11^.

A more rigorous understanding of the risk that 21^st^ century climate change poses to coral bleaching is needed to inform present-day management of Earth’s coral reefs and their ecosystems. Vital conservation strategies such as coral transplantation, genetic manipulation of corals, and engineering coral ecosystems can only be implemented at relatively small spatial scales^12,13^, meaning that they need to be prioritised to coral reef regions where future risk of coral bleaching is relatively lower. However, several methodological impediments have so far limited the ability to accurately pinpoint coral reef regions that are at lower risk of increased magnitude, duration and frequency of marine heatwaves. Firstly, most available projections of DHW derived from coupled atmosphere-ocean-general-circulation-models (AOGCMs) have been compiled from monthly-averaged rather than daily SST data (e.g., ^14,15^). This averaging masks important daily variation in temperature extremes, which accumulate as positive anomalies in the calculation of DHW^9^. Secondly, most future projections of DHW are currently available at a near-native resolution of AOGCMs (1° × 1°)^16–19^ (but also see^14,15,20^). Thirdly, existing forecasts of DHW do not account for inter-model variability from multiple AOGCMs using multi-model ensemble averaging, which has been shown to achieve better projection performance, regionally and globally^21^.

Here we introduce *CoralBleachRisk*, a continuous gridded (0.5° × 0.5° resolution) dataset of historic, current, and future coral bleaching risk based on downscaled and bias-corrected daily projections of SST and DHW. Downscaled and bias-corrected daily estimates are provided between 1985 and 2100 under three different future emissions scenarios from eight CMIP6 coupled AOGCMs. Annual summary projections of severity, duration, and onset of severe bleaching conditions are provided for three Shared Socio-economic Pathways (SSP) for the period 1985 to 2100. Projections are available for individual AOGCMs and a multi-model average. Tests show that historical projections of climate and bleaching closely reconstruct observed SST and past bleaching events, providing good confidence in future projections. We present two forms of data validation: (*i*) a climatological validation against satellite SST observations and (*ii*) an ecological validation against a historical dataset of past coral bleaching events. Our global projections of future risk of coral bleaching provide conservation managers and policy makers with the data needed to quantify 21^st^ century coral loss across the world’s coral reef regions.

## Methods

### Overview

An overview of the design of our dataset is provided in Fig. 1. Briefly, daily simulated sea-surface temperatures (SST) were available from eight Atmosphere-Ocean General Circulation Models (AOGCMs) and were retrieved from the open access Coupled Model Inter-comparison Project phase 6 (CMIP6)^22^ for the period 1985 to 2100 for three SSPs^23^. Data were downscaled to a 0.5° × 0.5° regular grid and bias-corrected against a high-resolution SST satellite dataset^24^. Downscaled and bias corrected daily SST data were used to calculate: (*i*) SST anomalies exceeding a spatially resolved bleaching threshold (defined as the MMM SST − a commonly employed bleaching threshold^8,9^); (*ii*) cumulative heat stress defined as degree heating weeks (DHW); (*iii*) and annual estimates of the onset, duration, and severity of coral bleaching conditions with DHW ≥ 4 or 8°C-week critical thresholds.

**Figure 1.**
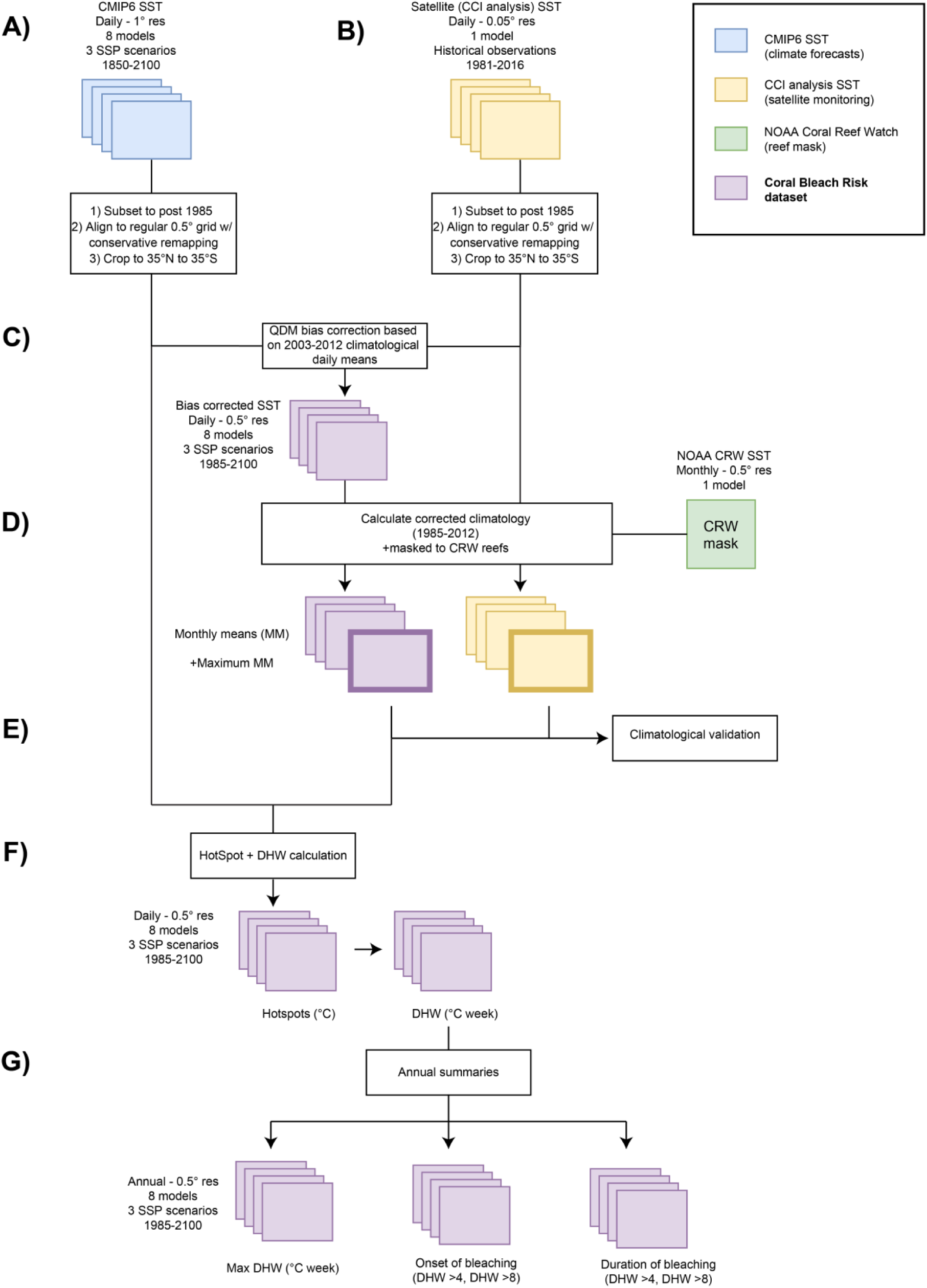
Processing steps and design of the CoralBleachRisk dataset. (**A**) Simulated projections of sea-surface temperatures (SST) from historical and future climates were extracted from 8 CMIP6 climate models for the period 1985 to 2100. (**B**) Satellite observations of SSTs (1985 to 2016) were extracted from the CCI analysis SST dataset. Datasets in A and B were re-gridded to a common 0.5° × 0.5° resolution. (**C**) CMIP6 projections were bias-corrected against the CCI analysis SST data using quantile delta mapping (QDM). (**D**) Climatological monthly mean SST were calculated from the downscaled and bias-corrected CMIP6 projections and CCI analysis SST data. (**E**) CMIP6 projections of mean monthly SST were validated against CCI analysis data from 1985 to 2014. (**F**) Hotspots (°C) and Degree Heating Weeks (DHW, °C-week) were calculated for each pixel at a daily resolution between 1985 and 2100. (**G**) Daily DHW data were used to calculate annual measures (onset, duration, severity) of severe bleaching risk in each pixel. These projections were validated against observations of coral bleaching for the period 1985-2010.

### Historical, future, and observed sea-surface temperature

#### Shared Socioeconomic Pathways

The latest AOGCM simulations of future climate are based on a combination of SSP and forcing levels used by the Representative Concentration Pathways (RCP) in CMIP5^25^. Three SSP scenarios were selected from hundreds of future climate scenarios currently available, to represent contrasting climate mitigation and adaptation challenges resulting in a wide range of different climatic futures^23,26^. Our three chosen scenarios represent: a ‘Middle of the road’, with medium socio-economic challenges for mitigation and adaptation (SSP 2-4.5); ‘Regional rivalries’, with high socio-economic challenges for mitigation and adaptation (SSP 3-7.0); and ‘Fossil-fueled development’, with high socio-economic challenges for mitigation and low socio-economic challenges for adaptation (SSP 5-8.5)^23^. We chose these because they reflect the most plausible future scenarios (i.e., SSP2-4.5 and beyond^27,28^). The SSP numerical designation for each simulation combines the socioeconomic scenario and radiative forcing level reached at the end of the century; e.g. SSP 5-8.5 is a fossil-fueled development scenario (SSP 5) with radiative forcing level reaching approximately 8.5 W/m^2^ by 2100^25,26^.

#### Historical and future temperature

Historical and future climate datasets with daily simulated SST available for each of the three SSP were extracted from the open access CMIP6 Earth System Grid Federation data portal (https://aims2.llnl.gov/search https://esgf-node.llnl.gov/search/cmip6/). Historical data (1850-2014) were accessed for 8 CMIP6 AOGCMs that also had daily SST available for SSP scenarios 2-4.5, 3-7.0 and 5-8.5^22^ (see Table 1 for model details). Future climate data were accessed for the period 2015-2100 for these same 8 CMIP6 AOGCMs^26^.

**Table 1:** Description of the eight AOGCMs used in our study. All Atmosphere-Ocean General Circulation Models (AOGCMs) had daily data available for the historical period (1850-2014) and three SSP scenarios (SSP2-4.5, SSP3-7.0, SSP5-8.5; 2015-2100). All models had a native nominal resolution of 1° × 1° and were re-gridded from their native non-regular grids to a regular 0.5° × 0.5° grid spanning 35°S to 35°N using conservative remapping. ECS = equilibrium climate sensitivity; change in global-mean air temperature due to an instantaneous doubling of CO_2_. TCR = transient climate response; warming from a simulation that is driven by an exponential 1.0 % per year increase in CO_2_. Models with a TCR inside the 1.4-2.2°C range (highlighted in green) are those included in the 5-model ensemble (*ens5*) as recommended in^52^.

| Model | Institution(s) | Ocean model | ECS | TCR |
| --- | --- | --- | --- | --- |
| *BCC-CSM2-MR* | Beijing Climate Center | MOM4-L40v2 | 3.02 | 1.59 |
| *CanESM5* | Canadian Centre for Climate Modelling and Analysis | NEMO-v3.4.1 | 5.64 | 2.66 |
| *CESM2* | National Center for Atmospheric Research | POP2 | 5.15 | 2.04 |
| *EC-Earth3* | EC-Earth consortium | NEMO-v3.6 | 4.26 | 2.38 |
| *IPSL-CM6A-LR* | L'Institut Pierre-Simon Laplace | NEMO-v3.6 | 4.70 | 2.32 |
| *MIROC6* | Center for Climate System Research, the University of Tokyo, the Japan Agency for Marine-Earth Science and Technology, the National Institute for Environmental Studies | COCO-v4.5 | 2.60 | 1.52 |
| *MRI-ESM2-0* | Meteorological Research Institute | MRI.COM-v4 | 3.13 | 1.56 |
| *NorESM2-MM* | Norwegian Climate Consortium | BLOM | 2.49 | 1.33 |

The period covering the historical simulations begins before significant anthropogenic climate change. Historic climatic simulations are forced by common observational datasets that account for changes in greenhouse gas concentrations from natural sources (e.g., due to solar variability and volcanic aerosols) and anthropogenic activities (e.g., CO_2_ concentration, aerosols, and land use)^22^. We used the historical climate simulations to test AOGCM performance against an observational dataset. The SSP scenarios simulate possible future climates from 2015, having been initialized using the climate conditions at the end of the historical period (2014) and the SSP simulations are effectively continuations of the historical simulations^26^. While there is no way to determine which of the model realizations should be preferred over others, and noting that each realization will lead to a slightly different climate state^29^, we opted to use only the first model realization (r1i1p1f1) for each AOGCM, which is common practice in climate change assessment tools (e.g., MAGICC/SCENGEN^30^). The rationale for this decision was that not all models had multiple realizations for the historical and future periods. Additionally, it has been established that projections of climate conditions improve with an increase in the number of ensemble-averaged models, not realizations^31^.

#### Observational data

Observed SST data were accessed from a high-resolution satellite derived dataset^24^, which was created as part of the European Space Agency Sea Surface Temperature Climate Change Initiative (CCI)^32^ (hereafter CCI analysis SST) available as an open access dataset at https://data.ceda.ac.uk/neodc/esacci/sst/data/CDR_v2/Analysis/L4/v2.1. We opted to use the level-4 (L4) gridded CCI analysis SST data described in Merchant, et al. ^24^. The L4 dataset is representative of global daily-mean SST (at 20cm depth) from 1981 to 2016 on a 0.05° × 0.05° grid, with gaps between available daily observations filled statistically. Validation of the L4 product using in situ measurements of SST from drifting buoys shows excellent spatiotemporal agreement, particularly after 2003^24^. A thorough product validation is provided in Rayner, et al. ^33^.

### Pre-processing of climate data

Different temporal extents and spatial resolutions of simulated and observed climate data required several pre-processing steps to ensure that the datasets were harmonised consistently, and that all data were on a spatially consistent grid (Fig. 1). All pre-processing was done using the Climate Data Operators (CDO) software^34^ and netCDF Operators (NCO) software^35^.

#### Re-gridding

To enable spatially consistent comparisons across all input datasets, the CMIP6 projections and the CCI analysis SST were re-gridded to a common 0.5° × 0.5° regular grid using conservative interpolation, an approach accounting for all source cells intersecting the destination cell and returning a weighted average^36^. Conservative interpolation was thus chosen to maintain areal averages between the coarse CMIP6 grids, which themselves were on different grid scales (Table 1), and the finer scale analysis grid^36^. This processing step was implemented with the *ncremap* function for NCO^35^. All re-gridded data were then cropped to latitudes between 35°N and 35°S where known coral reefs occur.

#### Temporal window

CMIP6 historical simulations start in 1850; however, only years since 1985 were included in *CoralBleachRisk* due to the modern observations of occurrence of mass bleaching events. For all datasets, any leap days (29^th^ February) were excluded from analysis as not all models contained leap days. To provide continuous estimates of SST between the historical and future periods, we harmonised the historic and forecast projections for each of the models by temporally merging past and future simulations^37,38^.

### Bias correction

Model bias correction is done to correct distributional biases in simulated climate data relative to historical climate observations^39^. We compared two-bias correction methods for our SST data: (*i*) a simple, commonly used delta change factor (CF) method, and (*ii*) the quantile delta mapping (QDM) method^40^. The CF method calculates anomalies between climatological averages (i.e., a baseline climate) of observed and simulated climatic conditions, and then adds those anomalies to the simulated climate. QDM is a multi-step bias correction method consisting of: (*i*) detrending the individual quantiles; (*ii*) applying a regular quantile mapping to the detrended series; and (*iii*) re-applying the projected trends to the bias-adjusted quantiles^40^. Regular quantile mapping involves mapping *n* quantiles of the cumulative distribution function of all values from the baseline period of the simulated climate onto the quantiles of the cumulative distribution function of all values from the baseline of the observed climate. For both methods, the baseline period − used to calculate climatological means for CF, and quantile calculations for QDM − was defined as 2003–2012 based on the high quality of satellite inputs to the CCI analysis SST during this period^24,33^ and a potential limitation in satellite-based representation of extreme temperature events prior to 2003^41^. The bias-corrected model SST values from each bias correction method were compared with the CCI analysis SST data for the period 1985–2014.

Bias correction methods are however unable to correct for systematic biases within a climate model (e.g., biases in circulation patterns, warming/cooling rates)^42^. Both methods assume that any biases in climate projections are stationary (i.e., that climatic characteristics in the baseline period will persist into the future), allowing alignment of the simulated baseline climate to the observed baseline climate. Because the QDM method explicitly preserves the change signal in all quantiles^40^ it is particularly suited to bias correcting climate data characterized by extremes and anomalies (e.g., rainfall^40^). Trend-preserving quantile correction methods have previously been demonstrated to provide the best downscaling methods for global climate models when attempting to preserve extreme values, and climate change signals^43,44^.

A statistical comparison of our two bias-correction methods was conducted over north-eastern Australia (140°E-160°E, 30°S-10°S) between 1985 and 2014. We chose this region due to the fact that CCI analysis SST data over the Australian region has been shown to be highly consistent with *in situ* tropical mooring measurements of SST, including for shallow reefs^33^; and because processing constraints prevent doing these tests over larger geographic extents. The comparison was done using Taylor diagrams that display correlations and standard deviations between the bias corrected and observed data, which confirmed that QDM method was superior to the CF method (see *Technical Validation*). Therefore, the QDM method was applied to all grid-cells across all years for each SSP, using the same baseline period.

### Climate metrics

We calculated six climate indices of coral bleaching risk for each reef location identified from the NOAA Coral Reef Watch dataset^45^ (Fig. 1). These heat stress metrics were: (*i*) Degree Heating Weeks (DHW, °C-week) – a cumulative metric of heat stress; (*ii*) annual maximum DHW; (*iii*) the 99^th^ percentile of DHW in each year; (*iv*) the annual number of days at or above DHW thresholds of 4 and 8 °C-week^9^. To calculate timing of onset of severe bleaching, we also calculated: (*v*) the day of year when DHW first exceeds 4 and 8 °C-week thresholds; and (*vi*) the day of year when DHW exceeds 4 and 8 °C-week, relative to the climatological coldest day of year in each coral reef pixel. The latter allowed global comparisons of the onset of bleaching conditions despite differences in the seasonality of SST between hemispheres. The calculation of each metric is detailed below.

#### Maximum Monthly Mean

The calculation of DHW requires defining the maximum monthly mean SST in each grid cell. This involved averaging daily (bias-corrected) SST data at each pixel, for each month over a 28-year period (1985-2012) to produce 336 (28 × 12) monthly mean SST values. To account for the climate drift in observation and projections of SST, we used linear regression to adjust the long term climatological monthly mean SST^46^. Specifically, a linear regression of the 28 yearly values for each month was calculated and solved at the mid-point of the years 1985-1990 and 1993 (which was the original climatology period used in satellite monitoring of heat stress^47^), thus giving a re-centered climatology. This technique was implemented to combine the benefit of a “long” climatology period whilst countering substantial modification to the historical baseline temperature that would result from averaging values across the 28-year period in which warming has occurred in reef locations^46^. Using the determined location-specific re-centered MMM climatology, we then calculated the Maximum Monthly Mean (MMM) as the maximum monthly SST for each pixel across the 12-month re-centered climatological period.

#### HotSpot metric and Degree Heating Weeks

Daily SST anomalies (i.e., HotSpots^9^) for day *i* (*HS_i_*, in °C) were calculated as the difference between *SST_i_* and MMM. *DHW_i_* (°C week) was then calculated as the cumulative sum of HotSpots above the MMM for each pixel over a rolling 12-week (84 day) window ending on day *i* as:

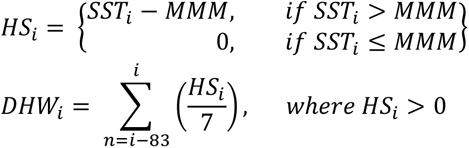

Traditionally, DHW values have been calculated using HotSpots of 1°C or greater as a high-pass filter^9,48^. However, recent work has shown an increase in the ability of statistical models to predict coral bleaching events using accumulations of all HS*_i_* > 0°C^49^. Moreover, the model-based bleaching outlook prediction tool produced by NOAA Coral Reef Watch incorporates a time-varying accumulation threshold for HotSpots that reduces from 1°C to zero within the first 10 weeks of the forecast period, in response to reduced variability in modelled temperature with lead time^50^.

#### Magnitude, duration, and onset of bleaching conditions

In each year from 1985 to 2100, we calculated the magnitude of coral bleaching risk as the annual maximum DHW in each pixel.

Given that exposure to summer-like temperature conditions is becoming longer, and the period of winter reprieve from warm temperatures is decreasing^51^, we calculated the duration of coral bleaching conditions as the total number of days, in a given year, when DHW exceeds the critical thresholds of 4 and 8°C-week. These thresholds have been associated with significant coral bleaching (i.e., Bleaching Alert Level 1) and widespread bleaching and coral mortality (i.e., Bleaching Alert Level 2), respectively^9^, and remain applicable when accumulating all HS*_i_* > 0°C^49^ (including within the bleaching outlook prediction tool^50^). While these associations were defined using ‘canonical’ DHW values calculated with the high-pass filter, recent bleaching observations have been reported at lower canonical-DHW values^1^. Regardless, the DHW methodology applied here and elsewhere is applied widely and consistent with the recent observations^9^.

We defined the onset of bleaching and severe bleaching conditions by the day number of each calendar year when DHW exceeds 4 and 8°C-week, respectively. To allow global comparisons of the onset of bleaching conditions that account for differences in seasonality between hemispheres, we also calculated relative onset of bleaching as the number of days since ‘mid-winter’ when DHW first exceeds 4 and 8°C-week. Here, mid-winter was defined as the climatological coldest day of the year for each pixel from the bias-corrected SST over the baseline period of 2003-2012.

### Multi-model ensemble

We generated two different multi-model ensemble averages based on different numbers of input datasets. The first ensemble (hereafter *ens5*) used five CMIP6 models that had a transient climate response (TCR; i.e. warming from a simulation that is driven by an exponential 1.0 % per year increase in CO_2_) inside the 1.4-2.2°C range as recommended by Hausfather et al.^52^ (Table 1). The second ensemble (hereafter *ens8*) was calculated using all eight models. Previous research using CMIP5 simulations has shown that the skill of multi-model ensemble averages to reproduce observed climate patterns asymptotes when at least five different models have been averaged^31^.

### Validation

#### Statistical validation of bias correction

Statistical validation of global climatological daily mean bias-corrected SST values from 1985-2014 for each individual model and our *ens5* hindcast was done by calculating their agreement with the CCI analysis SST data using the Kling-Gupta efficiency (KGE) metric^53^. Essentially, KGE ranges from –Infinity to 1, with values closer to 1 indicating more agreement between the bias-corrected and CCI analysis SST data. Each model’s daily and monthly climatological means, and those of *ens5*, were compared with the corresponding CCI analysis SST values based on the 2003-2012 period. Statistical validation was done for both bias-correction methods (QDM and CF). Taylor diagrams were used to assess pattern correlations and standard deviation ratios for our QDM bias-corrected SST estimates and for the CCI analysis SST at a global scale.

We also calculated multiple statistical metrics to quantify the relationship between our QDM bias-corrected SST estimates and the CCI analysis SST data for each of the 25 AR6 Working Group I reference regions^54^ and the 40 Marine Ecoregions of the World (MEOW)^55^ provinces. These metrics were: percentage bend correlation^56^, root mean square error, ratio of standard deviations, and modified index of agreement^57^. This statistical validation was first done using all cells within each of the IPCC AR6 reference and MEOW provinces and then repeated using only cells containing coral reefs according to the NOAA Coral Reef Watch dataset^45^.

#### Ecological validation

A separate validation was done to test the ability of our derived DHW metrics to predict the occurrence of past coral bleaching events^58^ within the IPCC AR6 regions. For this ecological validation, we used the open access global dataset of mass coral bleaching events collated by Donner, et al. ^58^ that spans most of the historical baseline period (1985 – 2012). This was the most comprehensive dataset available at the time of project development and analysis (available at https://figshare.com/projects/Coral_Bleaching_Database_V1/19753), noting that any more recent dataset (e.g., ^59^) would only have added 2 years to the historical baseline period (ending in 2014). Here we only considered records of severe bleaching events (i.e., severity level 3). This is because reports of lower or unknown bleaching severity can potentially conceal greater bleaching impacts if surveys occurred before or after the peak of bleaching – an important source of uncertainty in voluntary and citizen science programs^58,60^. This subset of the global mass coral bleaching dataset contained 1,975 severe coral bleaching events that overlapped with our historical projections of maximum annual DHW based on both the CMIP6 models and with the CCI analysis SST.

We quantified both the ability of DHW estimates from CMIP6 models and those calculated using CCI analysis SST data to predict the occurrence of observed severe bleaching events by calculating the hit rate (correctly projecting bleaching)^61^ based on the global mass coral bleaching dataset. According to this method, if a model predicted a coral bleaching alert (DHW ≥ 4°C-week) within 18 months of an observed bleaching event, this event was termed a “hit”. Alternatively, if no bleaching alert was predicted by the model (DHW < 4°C-week) within 18 months of an observed bleaching event, this event was termed a “miss”. These “hit” and “miss” occurrences were combined within the hit rate as follows:

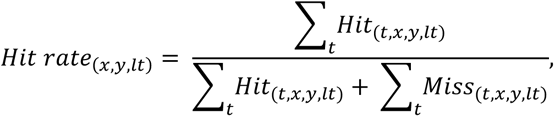

where *(x, y)* is the pixel containing at least one coral bleaching event, *t* is the survey date, and *lt* is the lead time for predicted bleaching alerts (18 months). The use of hit rate removes any skill bias by incorporating only correct positives (i.e., where a bleaching prediction matched an observed bleaching event^61^). We then calculated and mapped the mean hit rate and its standard error for each individual model, the five-model ensemble average *ens5*, and the CCI analysis SST within each IPCC AR6 region.

We used beta-regression^62^ to assess whether the proportion of severe bleaching events was well predicted by the *ens5* hindcast as well as the CCI analysis SST. To ensure sufficient sample sizes, the validation analysis excluded 7 (out of a total of 25 regions) IPCC regions that had < 10 severe bleaching records. The analysis was done using the rstanarm package in R^63^. Models were constructed with normal priors (*β_k_*~ *Normal*(0,2.5)) and four chains, each with 2000 samples, with the first 1000 samples being discarded as a burn-in. We ensured model convergence using Gelman-Rubin statistics (where values less than or equal to 1.1 were considered acceptable), along with testing for effective sample size, and visually examining trace-plots. We did posterior predictive checks to evaluate the model predictive accuracy relative to the observed data. We also plotted and compared the distribution of DHW values during severe bleaching events with that for all times within 1985—2012 at the same locations (i.e., baseline period) for both the *ens5* hindcast and the CCI analysis.

## Data Records

Access to our dataset is through Figshare (https://doi.org/10.25909/25143128). The gridded datasets are available as NetCDF files. The *ens8* data are not provided to limit the size of the online dataset, however, they can be recreated using CDO or NCO code provided.

The naming convention for the daily SST datasets is:

<*model*>_<*scenario*>_qdmCorrected_<*timeperiod*>.nc

where *model* is the name of the CMIP6 model, or “ens5” for the 5-model ensemble; *scenario* is the name of the scenario (historical, SSP2-4.5, SSP3-7.0, SSP5-8.5); and *timeperiod* is the temporal span of the file (e.g., 1985-2014). The monthly summary files share the same naming convention with an additional qualifier that identifies whether the values are representative of the minimum, mean, or maximum SST values seen within each month. All values are in °C.

The naming convention for the daily DHW is:

<*model*>_<*scenario*>_DHW_<*timeperiod*>.nc

where *model* is the name of the CMIP6 model, or “ens5” for the 5-model ensemble; *scenario* is the name of the scenario (SSP2-4.5, SSP3-7.0, SSP5-8.5); and *timeperiod* is the temporal span of the file (e.g., 1985-2100). Note that due to the nature of the DHW calculation the SSP outputs are each appended to the historical period (1985-2014).

The naming convention for the summary DHW files is:

<*model*>_<*scenario*>_<*timeperiod*>_DHWSummary.nc

where *model* is the name of the CMIP6 model, or or “ens5” for the 5-model ensemble; *scenario* is the name of the scenario (SSP2-4.5, SSP3-7.0, SSP5-8.5); and *timeperiod* is the temporal span of the file (e.g., 1985-2100). Note that due to the nature of the DHW calculation the SSP outputs are each appended to the historical period (1985-2014).

All gridded files have a 0.5° resolution with the following spatial dimensions – 140 latitude (35°S to 35°N) × 720 longitude (180°E to 180°W). Time periods differ for each file type. Daily SST data has *n* days where *n* is either 10,927 for the historical period (1985-2014) or 31,325 for the future SSP periods (2015-2100). The monthly SST summary files have *n* months, where *n* is either 360 for the historical period (30 years) or 1032 for the future SSP periods (86 years). The annual summary DHW time dimension is 115 years (1986 – 2100).

## Technical Validation

### Bias correction

CMIP6 projections were bias corrected using the quantile delta mapping (QDM) because testing showed that it better predicted observed SST on a subset of the study region — Australia’s Great Barrier Reef, which is the largest coral reef region in the world (Fig. 2A). Because of its ~14° latitudinal span, the Great Barrier Reef is known to display high levels of spatiotemporal variability in climate^64^, making it an excellent validation region. The strength of correlations with the CCI analysis SST using monthly areal averages for both bias-correction methods was comparable (QDM = 0.99 ± 0.003; CF = 0.97 ± 0.005), whilst the ratio of standard deviations was improved for the QDM method relative to the CF method, indicating less inter-model spread in variability (QDM = 1.01 ± 0.01, CF = 1.04 ± 0.05) (Fig. 2A). Significant improvements were shown for QDM over CF corrected datasets in the Kling-Gupta efficiency (KGE) metric when comparing individual model daily average SST with the CCI analysis SST (QDM = 0.91 ± 0.02, CF = 0.81 ± 0.07; *t*(7) = *3.725*, p = 0.007).

**Figure 2.**
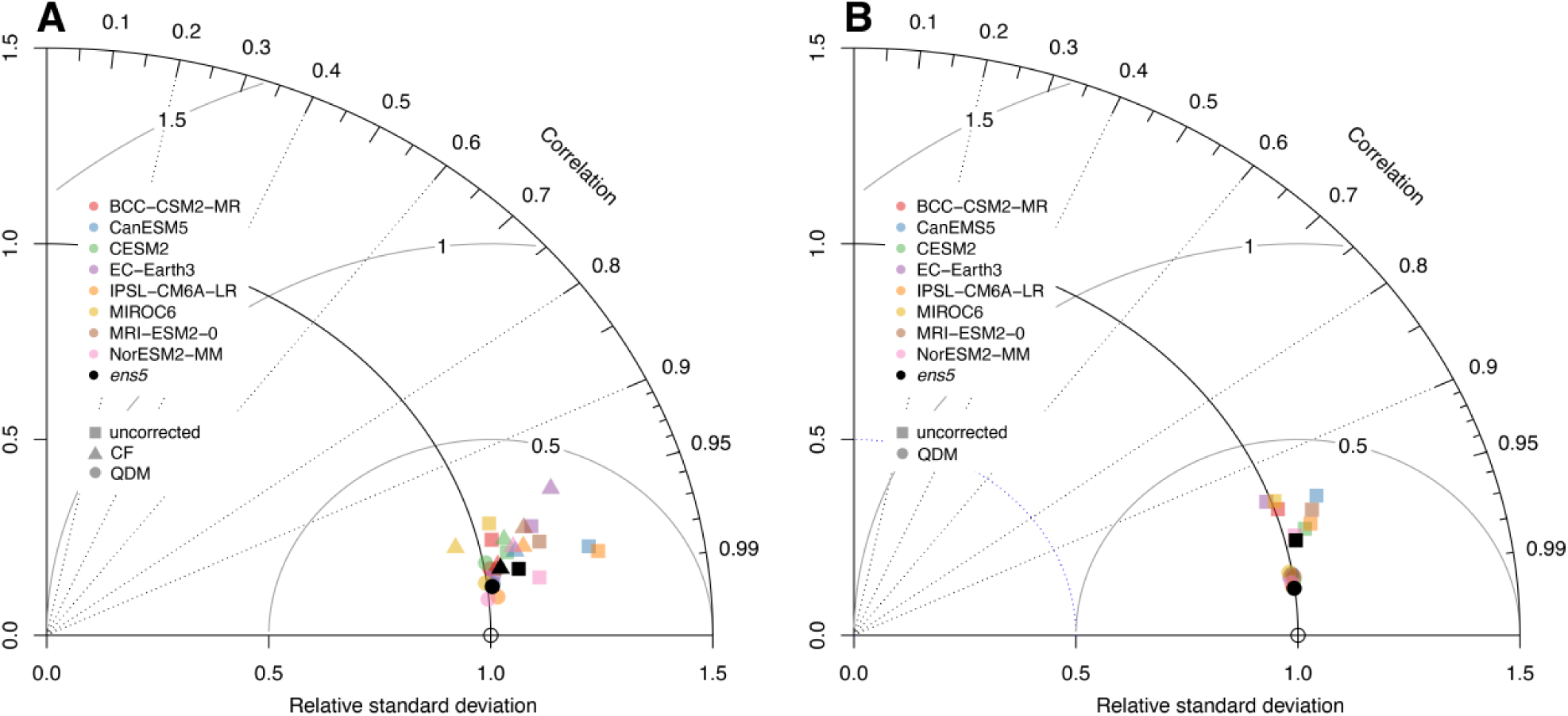
Taylor diagrams comparing projected and observed mean monthly SST. Comparison of mean monthly SST from eight CMIP6 climate models with satellite (CCI Analysis SST data) over the period 1985—2014 for (**A**) Australia’s Great Barrier Reef and (**B**) global coral reef regions. Uncorrected historical projections are shown as squares, change-factor (CF) bias corrected projections as triangles, and quantile-delta-mapping (QDM) bias corrected projections as circles. Five model ensembles (ens5) are shown for each of the datasets by the black dots. In both panels, colors represent different climate models (see inset key). A perfect model is one that corresponds to the empty black circle, i.e., with the same standard deviation as the CCI analysis SST data (i.e., relative standard deviation = 1), and a Pearson correlation coefficient of 1. Note that CF bias corrected projections are only shown for (A).

### Climate validation

At a global scale, our results demonstrate excellent agreement between monthly QDM bias-corrected CMIP6 SST data and the CCI analysis SST for the climatology period 1985-2014 (clustered circles in Fig. 2B). Our *ens5* hindcast was more skillful in capturing the observed SST conditions than any of the constituent models (Fig. 2B). Statistical validation of monthly climatological means for IPCC AR6 regions showed similar patterns (Fig. 3), with average monthly percentage bend correlations of 0.91 (S.D. = 0.10). The *ens5* hindcast did better than the constituent models with an average monthly percentage bend correlation of 0.93 (S.D. = 0.07). Restricting the comparison to reef cells only decreased the average correlation across models and regions slightly to 0.86 (S.D. = 0.13), while for the *ens5* hindcast it reduced the bend correlation to 0.89 (S.D. = 0.09). Similar patterns were observed for RMSE with average values of 0.50 (S.D. = 0.23), and an *ens5* average of 0.42 (S.D. = 0.17) across all cells. RMSE values decreased when looking at reef cells only, with average values of 0.44 (S.D. = 0.19), and an *ens5* average of 0.38 (S.D. = 0.17). KGE statistics were between 0.51 and 0.77 depending on the model considered (mean = 0.68, S.D. = 0.08). The strength of the KGE statistic improved when using *ens5* (KGE = 0.81). The range of KGE statistic for all AOGCMs narrowed when using projections of SST for reef cells only, with values between 0.57 and 0.77; and the mean was marginally lower (mean = 0.66, S.D. = 0.09). Likewise, the KGE statistic decreased for the *ens5* hindcast for reef cells only (KGE = 0.73) compared with when including all cells.

**Figure 3:**
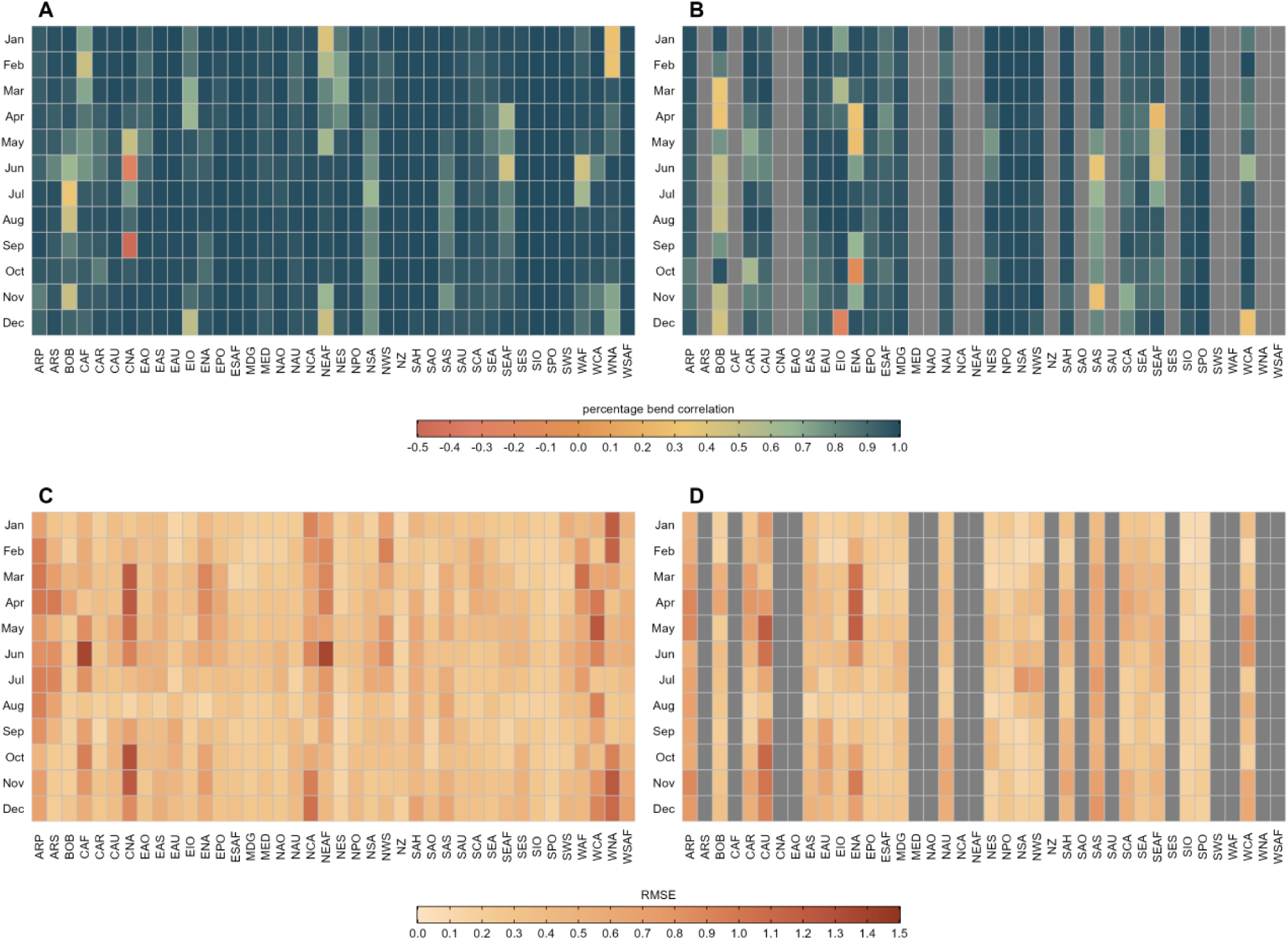
Validation of SST values in IPCC AR6 reference regions. Comparison of QDM bias-corrected CMIP6 projections of monthly SST with CCI analysis SST data for IPCC AR6 reference regions for the period 1985-2014. Results are shown for a five-model ensemble (*ens5*) projection. Upper panels show the percentage bend correlation (**A**) for all pixels and (**B**) reef cells only. Lower panels show the root mean square error (RMSE) for (**C**) all pixels and (**D**) reef cells only. Grey colour in (**B**) and (**D**) indicates no reef cells present in that specific region. With ARP: Arabian Peninsula; ARS: Arabian Sea; BOB: Bay of Bengal; CAF: Central Africa; CAR: Caribbean; CAU: C Australia; CNA: C North America; EAO: Equatorial Atlantic Ocean; EAS: E Asia; EAU: E Australia; EIO: Equatorial Indian Ocean; ENA: E North America; EPO: Equatorial Pacific Ocean; ESAF: E Southern Africa; MDG: Madagascar; MED: Mediterranean; NAO: N Atlantic Ocean; NAU: N Australia; NCA: N Central America; NEAF: N Eastern Africa; NES: NE South America; NPO: N Pacific Ocean; NSA: N South America; N W South America; NZ: New Zealand; SAH: Sahara; SAO: S Atlantic Ocean; SAS: S Asia; SAU: S Australia; SCA: S Central America; SEA: S E Asia; SEAF: S Eastern Africa; SES: S E South America; SIO: S Indian Ocean; SPO: S Pacific Ocean; SWS: S W South America; WAF: Western Africa; WCA: W C Asia; WNA: W North America; WSAF: W Southern Africa.

Similar patterns were seen in the Marine Ecoregions of the World (MEOW) provinces, with an all-model-average monthly correlation of 0.86 (S.D. = 0.14) for the 163 MEOW provinces in our latitudinal range. This correlation increased for the *ens5* hindcast to 0.89 (S.D. = 0.10). RMSE values followed similar patterns with mean RMSE across all models of 0.43 (S.D. = 0.25) compared with 0.35 for *ens5* (S.D. = 0.18). Masking to reef cells decreased the average correlation to 0.81 (S.D. = 0.15) and 0.83 (S.D. = 0.13) for all models and the *ens5* hindcast, respectively. RMSE values improved when considering reef cells only, with average values of 0.35 (S.D. = 0.16) and 0.29 (S.D. = 0.12), respectively. The full statistical validation for the IPCC AR6 regions and the MEOW provinces is available in Supporting Data 1.

### Ecological validation

Tests of projections of bleaching events against an existing dataset of coral bleaching^58^ show that both the *ens5* hindcast and the CCI analysis SST predict bleaching events with high accuracy, with hit rates > 80% in 22 and 21 regions out of a total of 26 regions, respectively (Fig. 4). Bayesian beta-regression modelling indicates no significant differences between the *ens5* hindcast and the CCI analysis SST data in their capacity to predict observed bleaching events in the 18 regions with > 10 bleaching events between 1985 and 2012. This high predictive capacity was not affected by the variable number of bleaching events across regions for either dataset (*ens5* = 0.05, 95% CI −0.31:0.46; CCI analysis SST = −0.07 95% CI −0.41:0.33).

**Figure 4:**
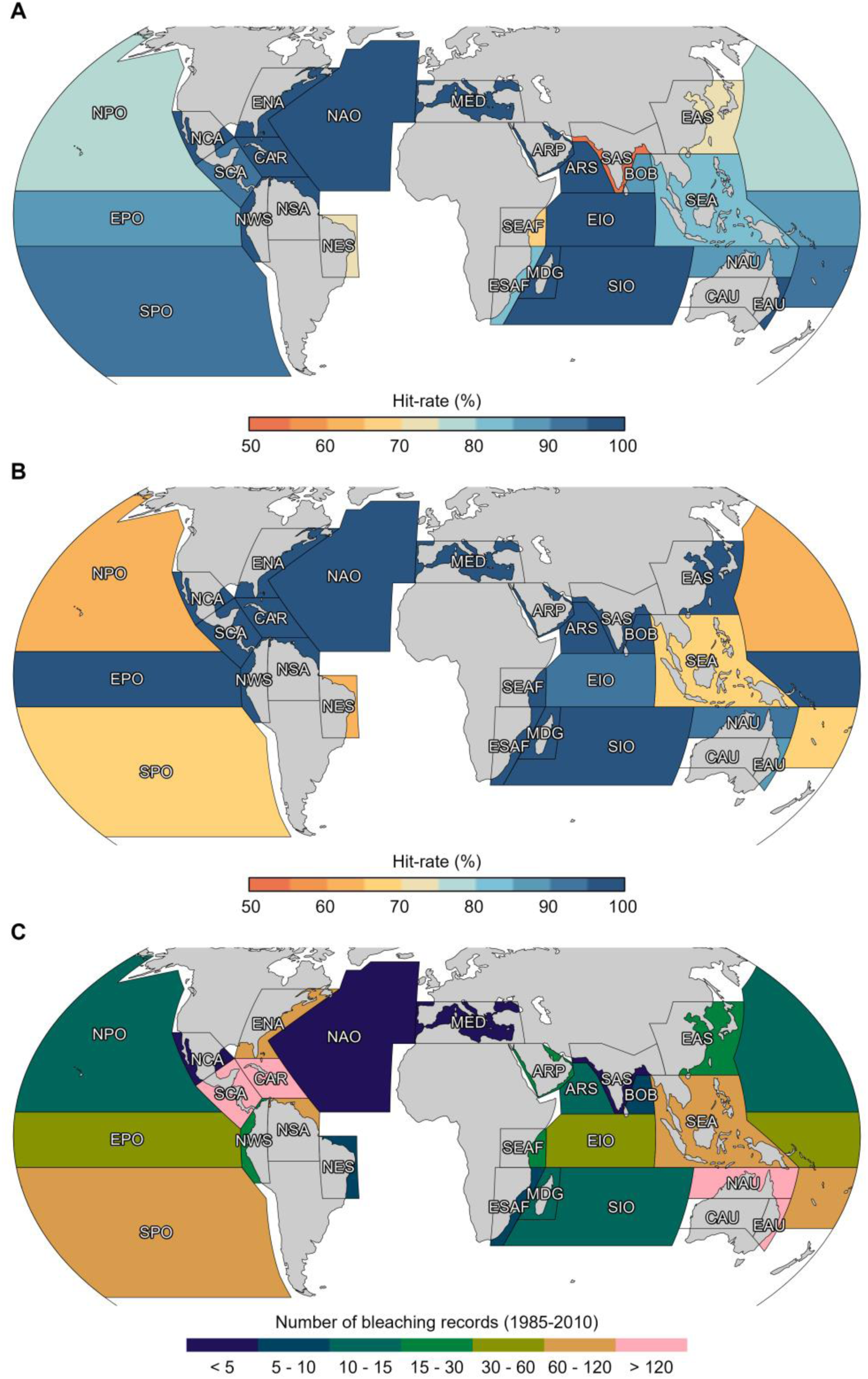
Comparison of projected and observed bleaching events in IPCC AR6 regions. Percentage of successful bleaching events predicted (hit rate) between 1985 and 2012 for (**A**) the five-model ensemble (*ens5*) and (**B**) the CCI analysis SST data. (**C**) The number of severe bleaching events (total = 1,975) recorded in each IPCC AR6 region. Details of IPCC AR6 region codes are given in the Figure 3 caption.

A region-level analysis indicated that *ens5* hindcast was better able to recreate the number of individual bleaching events than the CCI analysis SST data in 5 of the 18 regions that had >10 bleaching events recorded (Caribbean *X^2^*(1) = 5.88, p = 0.015; South East Asia *X^2^*(1) = 3.89, p = 0.048; Southern Pacific Ocean *X^2^*(1) = 18.58, p < 0.001; Eastern Australia *X^2^*(1) = 11.43, p < 0.001; South East Africa *X^2^*(1) = 4.16, p < 0.04) (Fig. 4). However, when comparing across all regions (i.e., globally), there was no statistical support that the *ens5* hindcast could more, or less, accurately predict the total number of regional scale bleaching events when compared to the CCI analysis SST data (*V* = 56, p = 0.552). We found a mean hit rate of 90% (S.D. = 13%) across the IPCC AR6 regions the for *ens5* hindcast (Fig. 4A), compared to a mean hit rate of 92% (S.D = 13%) from CCI analysis SST data (Fig. 4B). All posterior model checks, including the Gelman-Rubin statistic, effective sample size, and traceplots, indicated that the models were constructed correctly and showed no evidence of lack of convergence or autocorrelation.

As with the statistical validation, the *ens5* model had improved performance over the constituent individual models in all IPCC AR6 regions (Fig. 5A). Individual model skill varied greatly between regions, with some regions such as the North Pacific Ocean (NPO) and South Pacific Ocean (SPO) showing reduced predictive skills for all models, and the satellite monitoring CCI analysis SST data, as well as greater hit rate variability (Fig. 5B). DHW values during the baseline period showed similar distributions (mostly < 1°C-week) for the *ens5* hindcast and the CCI analysis SST. In each case they were distinct from the distribution of DHW values during severe bleaching events, suggesting that both datasets enable the distinction of baseline from bleaching conditions (Fig. 6).

**Figure 5:**
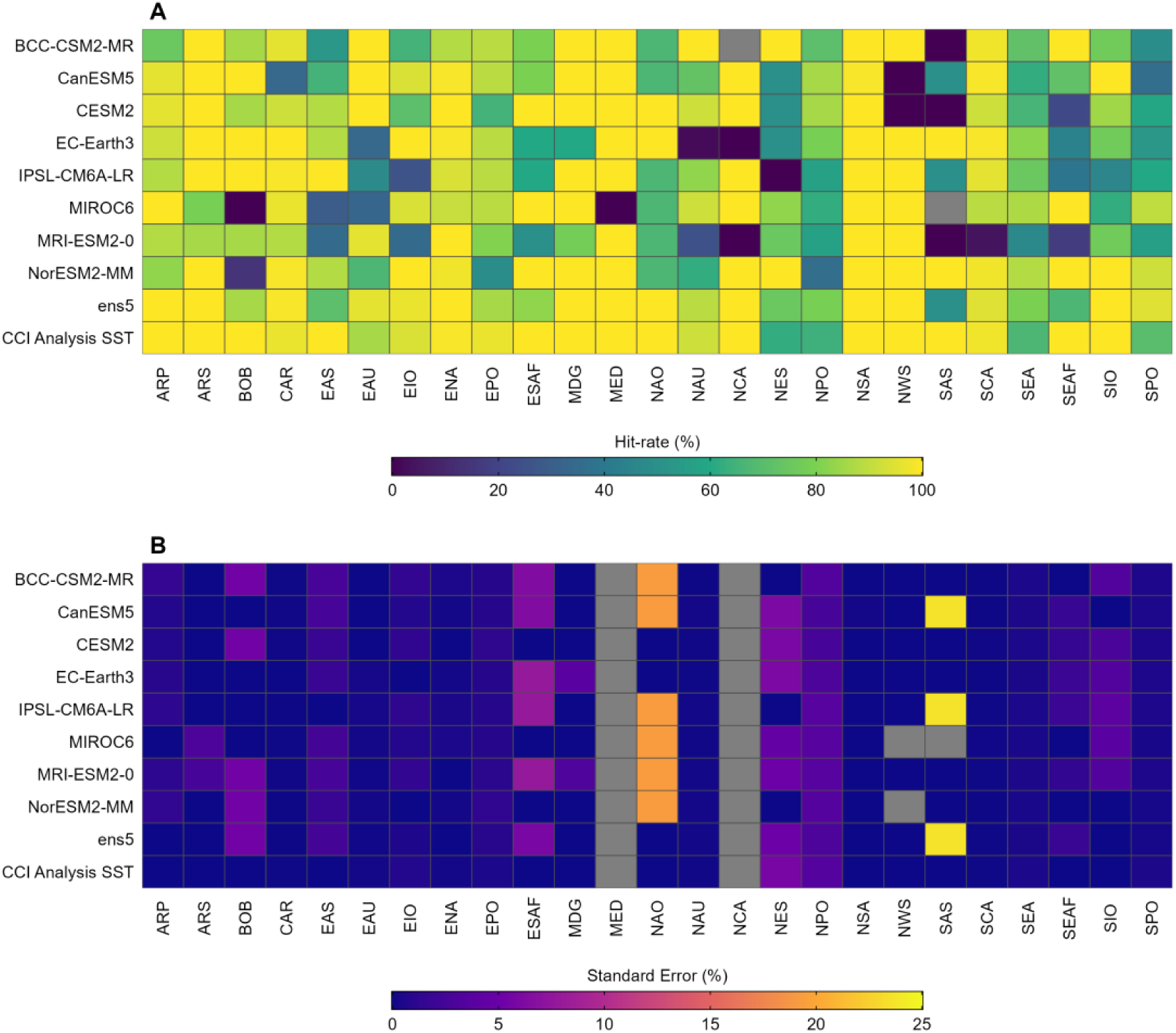
Comparison of projected and observed bleaching events for individual climate models. Assessment of the ability of the downscaled and bias-corrected CMIP6 models to effectively predict observed coral bleaching events in IPCC AR6 regions measured as (**A**) mean % hit rate and (**B**) % hit rate standard error (S.E.). Grey colour for a given model × region combination indicates no data due to insufficient bleaching records. CMIP6 models (individual models and five-model average *ens5*) and the CCI analysis SST are shown as rows, with IPCC AR6 regions shown as columns (see Fig. 3 for a description of region names and Fig. 4 for region locations).

**Figure 6:**
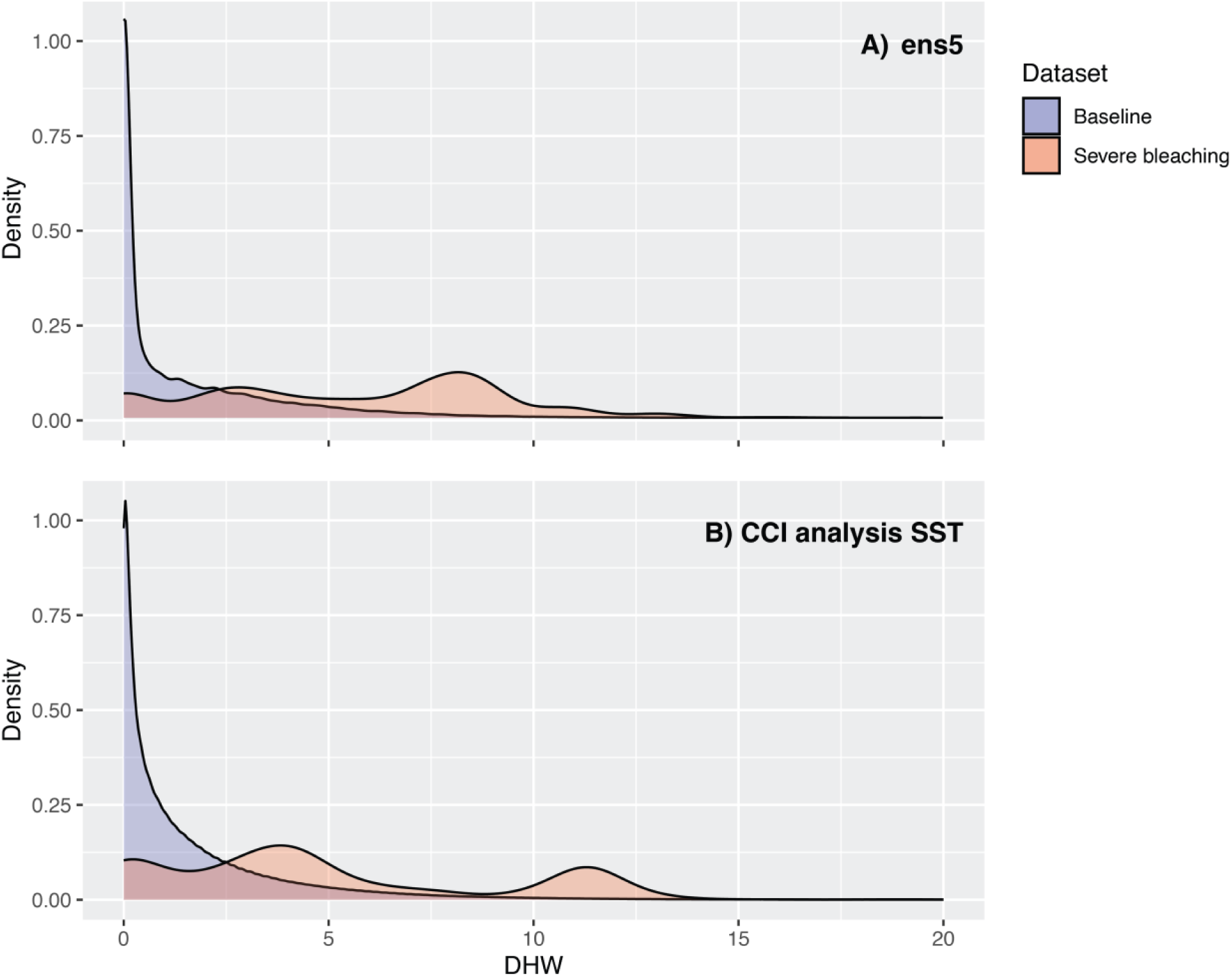
Distribution of DHW during severe bleaching events and throughout the climatological baseline period. Density plots show DHW distributions only during severe bleaching events within 1985-2012 (red) and throughout the baseline period (i.e., all days considered between 1985-2012 at the same locations; blue) for (**A**) the *ens5* multimodel ensemble and (**B**) the CCI analysis SST.

## Usage Notes

*CoralBleachRisk* provides access to daily projections (1985-2100) of SST (in addition to monthly minima, maxima and means) from eight CMIP6 models. Moreover, it provides validated hindcasts and future projections of daily, monthly, and annual estimates of risk of bleaching for Earth’s coral reefs. These include HotSpots^9^ above the maximum monthly mean SST (i.e., the location-specific expected summertime maximum temperature) and resulting daily DHW between 1985-2100 for a total of > 74,500 0.5° grid-cells, containing ~1500 reef locations (Fig. 1). Annual summary metrics of coral bleaching risk derived from this daily data include minimum, maximum, mean, and standard deviation of DHW, the onset and duration of DHW above the 4 and 8-°C week thresholds. All data are accessible through FigShare at https://doi.org/10.25909/25143128.

An accompanying online *CoralBleachRisk* data portal (https://coralbleachrisk.net/) allows users to graphically explore, download, and analyse the timing, duration, and extent of the coral bleaching risk for different years, scenarios, and climate models (Fig. 7). Data can be plotted and downloaded for a specific region (user-defined, IPCC reference regions^54^ or Marine Ecoregions of the World^55^ [MEOW] provinces) or locations with individual reefs, using the data portal. Available metrics include annual summaries of coral bleaching risk, specifically minimum, maximum, mean, and standard deviation of DHW, the onset of DHW above the 4- and the 8-°C-week threshold, and the duration of DHW above each threshold. Graphical assessments of the changes in annual maximum DHW, the annual onset of bleaching conditions, and the annual duration of bleaching conditions are also provided. A short tutorial is available in the data portal explaining how to crop, extract, and plot multiple metrics of SST and coral bleaching risk from different climate models.

**Figure 7:**
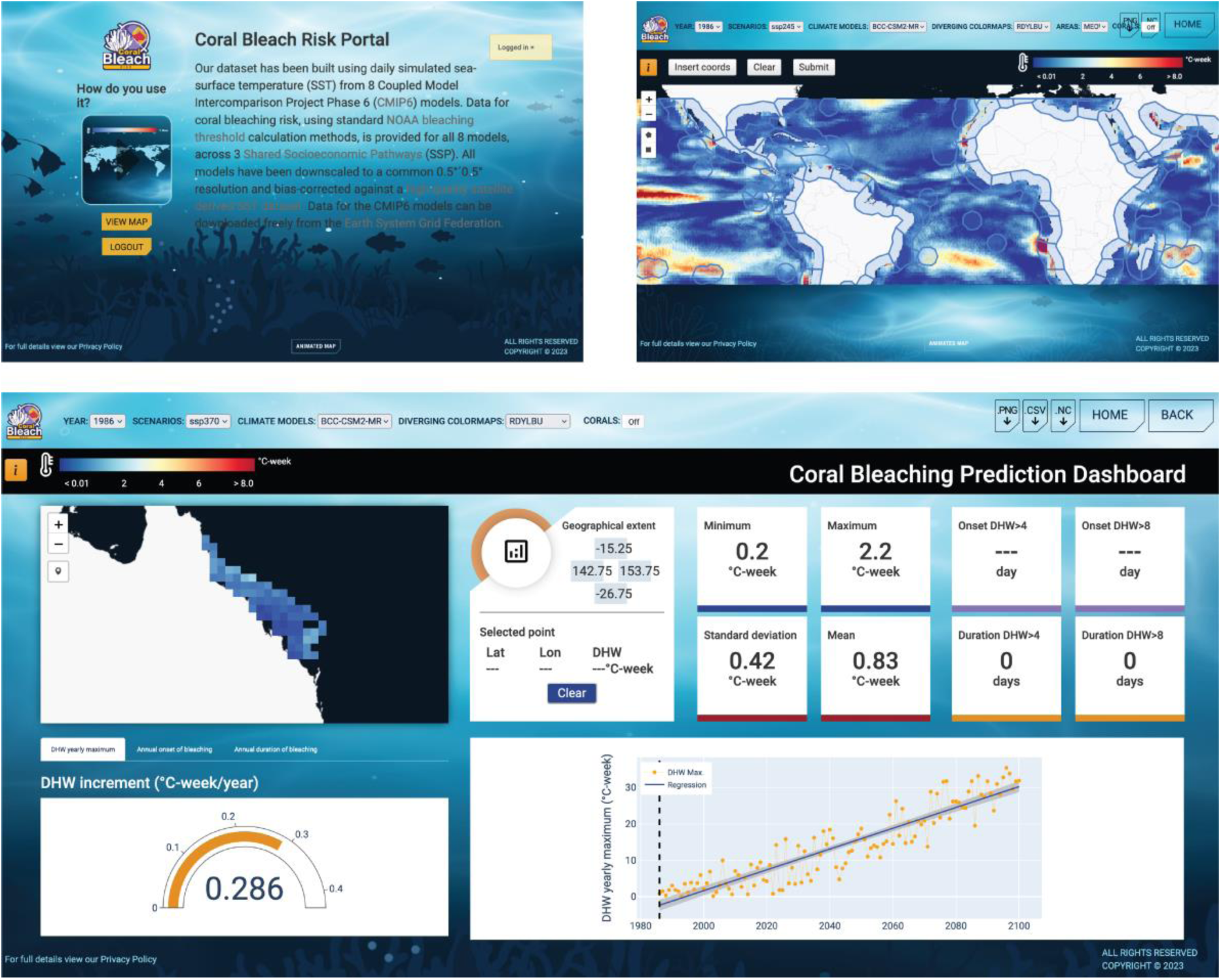
Snapshot of the *CoralBleachRisk* online data portal. Portal is available at https://coralbleachrisk.net/.

*CoralBleachRisk* provides a data framework and supporting code that can be updated and built on. Future advances could include incorporating additional climate models, scenarios and realizations into *CoralBleachRisk*^65^. However, this comes with the potential caveat that the number of models for analysis might need to be reduced. When CoralBleachRisk was developed, there were only sufficient CMIP6 climate model data to compare the first realization of three SSP scenarios for a total of eight models. Moreover, only five of these models (used in the *ens5* projection) had suitable ‘best’ estimates of climate sensitivity^52^. The size of this model ensemble projection is not problematic, because the performance of multi-model averaged climate projections tend to plateau with five or more models^31^. Likewise, our use of a single realization in ensemble projections is unlikely to affect accuracy because model performance increases to this asymptote faster with the addition of more models rather than more realizations^31^. Further methodological improvements could include refining the spatial resolution of *CoralBleachRisk* projections using dynamic downscaling^15^. However, this could only be done regionally, because it is not yet computationally feasible at a global scale. Lastly, as global temperatures continue to rise above maximum summertime averages, some level of coral adaptation is likely to occur over decadal time scales^19^, meaning that the relevance of traditional methods for calculating DHW and associated bleaching thresholds might need to be re-evaluated.

All analyses were performed in Climate Data Operators (CDO) software^34^ and netCDF Operators (NCO) software^35^, and R version 4.4.4^66^ using packages terra 1.6-48^67^, sf 1.0-9^68^, loadeR^69^, rnaturalearth^70^, plotrix^71^ and the climate4R package bundle^72^.

## Code Availability

Code used to bias correct and generate the metrics presented in this paper is available at https://doi.org/10.25909/25143128.

## Acknowledgements

The authors wish to acknowledge funding from the following sources: Australian Research Council grants FT200100870 (CM) and DP230102986 (SFH), SUBAK Fellowship grant (SCB), and seed funding from University of Adelaide’s Environment Institute.

## Author contributions

Conceptualization: CM, SCB, DAF

Methodology: CM, SCB, DAF, SFH

Investigation: CM, SCB

Visualization: CM, SCB

Supervision: CM, DAF

Writing—original draft: CM, SCB

Writing—review & editing: CM, SCB, DAF, SFH

